# Novel algorithms deriving clinical performance measures from smartphone sensor data collected under a walking test

**DOI:** 10.1101/2021.10.21.465337

**Authors:** Max A. Little, Sami Volotinen, Brad Sanderson, Ulla Huopaniemi, Florence Mowlem, Jennifer Olt, Bill Byrom

## Abstract

The modern ubiquity of smartphones which incorporate sensors such as inertial measurement units (IMU) and global positioning system (GPS) receivers, raises the possibility that they might be used as inexpensive devices for clinical performance testing. Here, we report on the development and testing of an app and associated statistical signal processing algorithms, to measure the basic gait properties of step rate, walking speed and distance under an outside short walk test experimental protocol, from digital accelerometry and GPS. We provide extensive details on the assumptions underlying the development of the algorithms. A small set of experiments across a range of conditions and individuals, demonstrate the reliability of the combined app, protocol and algorithms. With reasonable adherence to the test protocol, these experiments show 2% error across all metrics. The emphasis in this paper is on illustrating the design principles which all such systems, aiming to re-purpose consumer smartphones as clinical gait measurement devices, must address.

## 1 Introduction

Clinical drug development programs aim to gain sufficient insights into intervention effects to enable effective investment decision making, and to support ultimate regulatory submission and drug labeling. The cost of drug development is estimated to be around US$2.6 billion [DiMasi et al., 2016] and the rate of candidate termination during drug development is reported to be 81%, with a 55% attrition rate between phases II and III [Kaitin and DiMasi, 2011]. Early no-go decision making can lead to significant savings.

Today’s clinical trials are employing greater digitalization and leveraging more at-home assessments to improve the quality and reliability of data, enable richer and more frequent measurement of health outcomes, and make study participation more convenient by reducing the frequency of in-clinic visits. Smartphones are regularly used in clinical trials to provide access to apps to collect patient-reported outcome measures (PROMs), provide engagement solutions to guide patients throughout the study, and provide interaction and connectivity with the study site. Smartphones also afford the opportunity to collect objective measurements from patients, by leveraging their on-board sensors in novel ways not originally intended. For example, there is huge growth in the number of apps available on both Apple (Apple Inc., Cupertino, CA, USA) and Google (Alphabet Inc., Mountain View, CA, USA) platforms focused on personal health and fitness [Middelweerd et al., 2014], many of which leverage the inbuilt sensors (such as the accelerometer and GPS) to provide valuable features such as activity tracking.

Leveraging smartphone sensors to develop new clinical endpoint measures affords the possibility to measure things that were previously difficult or impossible to measure, to measure more frequently by enabling assessment from home rather than during clinic visits, and to study elected patient functioning in real-world conditions that may compliment established in-clinic tests. Smartphone apps accessing the inbuilt sensors may enable this to be done conveniently and simply without the need to supply an additional peripheral device such as a wearable. Because, unlike wearables, smartphones are not always carried, they are less suited to continuous monitoring and providing accurate daily totals or summaries. They are, however, better suited to the instrumentation of sensor-enabled discrete short performance outcome assessments.

Performance outcome assessments are a type of clinical outcome assessment, and are defined as “a measurement based on standardized task(s) performed by a patient that is administered and evaluated by an appropriately trained individual or is independently completed” [Cagney et al., 2017]. Examples include completing a timed 25-foot walk test, or a memory task such as a word recall test. Large observational studies have illustrated the potential of using smartphone components to collect performance outcome measures. For example, the MyHeart Counts Cardiovascular Health Study [McConnell et al., 2017] collected motion data from over 20,000 adult participants across the US, with almost 5,000 also completing an app-delivered six-minute walking test, with all data collected using an app developed using Apple ResearchKit [Mohammadi, 2015].

Pharmaceutical companies have also driven innovation in this area. Roche (F. Hoffmann-La Roche Ltd., Basel, Switzerland), for example, developed a set of smartphone-delivered performance outcome assessments for use in the study of treatments for Parkinson’s disease [Lipsmeier et al., 2018]. Tasks include a phonation test to measure voice degeneration, simple tests of balance and gait using the accelerometer to measure sway and stepping, a finger tapping dexterity test using the smartphone touchscreen, and tests to measure tremor using the device accelerometer while the patient held their smartphone with arm extended for 30 seconds. GlaxoSmithKline (London, UK) included a measure of wrist range of motion in their PARADE study of rheumatoid arthritis using the smartphone gyroscope and accelerometer sensors to measure wrist mobility while the patient performed a number of defined movements [Crouthamel et al., 2018, Hamy et al., 2020].

Smartphone technology provides a convenient approach to implement, collect and transmit sensor data for performance tests, but additional work is required to develop and validate algorithms to interpret the data collected and translate these data into clinical outcome measures. Walking-based performance tests typically require the estimation of distance walked or aspects of gait such as the number of steps taken. While the use of accelerometer data to estimate stepping events during walking is not novel [Bassett et al., 2017], there are many approaches to the measurement of steps and associated gait parameters from 3D raw acceleration data. In addition, using sensor data from a smartphone to estimate steps and distance brings additional challenges to the algorithm developer as the device was not built for this purpose and this can introduce additional challenges and variability into the data that impact estimation. For example, unlike a wearable sensor the smartphone unit may not be firmly affixed to the body and this may introduce additional noise in the acceleration signal due to extraneous and sympathetic movement of the unit while walking, and less standardization in placement location and orientation may introduce additional variation between uses.

In this paper, we describe the development and pilot performance of algorithms to measure step rate, walking speed and distance walked using GPS and accelerometer data sampled from the on-board smartphone sensors using a proprietary app during the conduct of a 6-minute short walking test intended to be conducted outside. Since smartphone sensor-based testing is still quite novel, our aim is to make the relevant technical issues as clear as possible with the hope of opening up further research in this promising area. Thus, we emphasise that the primary aim of this paper is to describe the technical issues we encountered and how they can be addressed, noting that this should not be misunderstood as advocating for the *only* or ultimately optimal, way in which such issues can be solved. Furthermore, the validation in the paper is from a very small cohort of self-experimenters, it should therefore not be misinterpreted as representing a fully rigorous algorithm validation study. We discuss the implications for these limitations below.

## 2 Algorithm development

The problem of tracking walking behaviours using physical motion sensing devices, has a long history. Perhaps the oldest are *pedometers* (step counters) and *odometers* (typically, distance wheels) which started out as specialized mechanical devices and are these days largely replaced by *microelectromechnical inertial measurement units* (IMUs) and *global positioning systems* (GPS) and specialized software algorithms. These are sufficiently miniaturized to fit inside today’s smartphones and smartwatches. The flexibility of such hardware/software combination raises the possibility of measuring much more than just step counts and stride lengths, including quantifying changes to multiple, subtle aspects of gait performance for medical applications. To be concise we will restrict discussion to step rate, speed and distance estimation.

### 2.1 Controlling for test protocol

Because human behaviour is, in general, inherently unpredictable, developing algorithms to process such data either requires establishing the environmental and/or behavioural context of the measurements, or requires specifying the conditions under which such an algorithm will function correctly. This is essential to avoid the worst effects of *ambiguity*, *confounding* and *collider bias* [Pearl, 2009]. An ambiguous behaviour is one which, without further establishing context through other forms of physical measurement, cannot be discriminated in principle from other, irrelevant behaviours. For example, from IMU measurements, walking on a treadmill may be indistinguishable from walking outdoors. Similarly, from the point of view of GPS measurement, traveling slowly on a bicycle may be indistinguishable from running. Therefore, to quantify, for instance, stride length, it is clear that neither GPS nor IMU measurements alone may suffice. In principle, it may be possible to *fuse* GPS and IMU measurements in order to reduce such ambiguities, but there will always be some ineliminable level of behavioural ambiguity.

Similarly, confounding occurs when a target measurement has one or more behavioural and/or environmental factors which cause ambiguities, but additionally, there are causal relationships between these factors themselves. In which case, it may be impossible in principle to use sensor device fusion alone to solve the problem. For example, age is a major factor in determining normal stride length, but various diseases of aging affect gait. Thus, it may be impossible to detect the gait-dependent existence of a specific disease without also knowing the age of the participant and incorporating this as part of the algorithm. This may mean that age-independent measurement of certain gait characteristics is impossible under certain circumstances.

Considering multiple causal factors and relationships in gait measurement, there are additional paradoxical phenomena such as *collider* (in particular, *selection*) *bias*. To get around many of these problems, experimentalists use (diagnostic, randomized) *controlled trials* (RCTs) in order to isolate behavourial and/or environmental causes and their measurement effects [Matthews, 2006]. Intuitively, establishing control over behavioural and/or environmental factors, in principle allows the measurements to be attributed to those factors alone. For example, by controlling for walking outside the home, we clearly rule out the ambiguity that individuals may be walking on a treadmill indoors.

Enforcing such control under supervised lab conditions is straightforward, but the goal of remote monitoring is to do this unsupervised, outside the lab. One common approach is to set up software on a smartphone which prompts the user to follow some kind of test protocol. In that way, provided the user follows the prompts, it is possible to simulate the experimentally-controlled condition remotely, referred to as a performance outcome test. Of course, it is not always possible to follow prompts remotely, but it is sometimes possible to quantify adherence algorithmically, although this is not a perfect solution [Badawy et al., 2018]. In this study, for simplicity, we will assume that adherence to test protocol is sufficiently reliable, recognizing that this is a problem which will need to be addressed in practice.

### 2.2 Pre-processing inertial sensor data

MEMS IMUs generally come equipped with three different sensor devices: accelerometers, gyroscopes and magnetometers. For our purposes, we will focus on accelerometry as this will suffice for quantifying gait performance. Pre-processing this data requires taking into account the basic physical kinetics involved. The IMU device having negligible mass itself, is attached to the relatively much heavier, smartphone of fixed mass. By Galilean invariance, the smartphone experiences the vector sum of all individual forces acting on it and according to Newton’s 2nd law, the acceleration measured by the IMU is proportional to the sum of these forces. Further, by Einstein’s inertial equivalence principle, one of the forces in this sum is any effective “reaction” force the smartphone experiences which prevents it from free-falling towards the centre of the Earth under gravitational acceleration. The magnitude of this force will therefore be zero if the smartphone is in free-fall. This reaction acceleration points away from the center of the Earth and effectively “orients” the IMU.

In this application, this “gravitational reaction force” is of no interest to measuring aspects of the behaviour of the user, because we expect that the smartphone is tightly mechanically coupled to the user, and the user themselves is not falling down, either. However, it may be useful for determining the “orientation” of the smartphone relative to the user as we assume the user is upright. For example, in our test protocol, the smartphone is contained in a waist bag at the small of the back. While orientation must be landscape, there remains four possible ways in which the smartphone can be oriented. Determining this vector is necessary, therefore. Let us formalize the problem as one of determining the gravitational acceleration vector 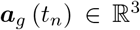 at any measurement time instant *t_n_* for *n* = 1,…, *N*, from the vector measured by the IMU 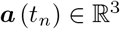:

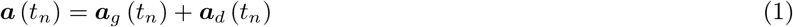

where 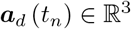 is the “dynamic” acceleration due to the movement of the smartphone user alone. Clearly, this problem can only be solved given further information. In particular, if the dynamic acceleration has zero magnitude ∥***a**_d_* (*t_n_*)∥ = 0 then ***a*** (*t_n_* = ***a**_g_* (*t_n_*). This offers an opportunity to “calibrate” smartphone orientation as long as the user does not move. This is not practical in our application, however, so we need to determine ***a**_g_* (*t_n_*) by making additional assumptions, in particular, that either:

1. orientation remains fixed throughout the test, or,
2. orientation changes on a time scale which is large by comparison to the timescale of changes in dynamic acceleration; for example, a device in a slightly loose bag worn by the user rotates slowly over the duration of the test.

We will discuss both assumptions next.

#### 2.2.1 Fixed orientation

Under assumption (1) above, the time *t* of measurements is irrelevant in which case ***a*** (*t_n_*) = ***a**_g_* + ***a**_d_* (*t_n_*). If we assume that **a**_d_ behaves like some random “disturbance”, an appropriate summary statistic of the distribution of ***a*** (*t_n_*) would suffice as an estimate of ***a**_g_*. In particular, if we assume that 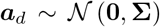 is zero mean, Gaussian distributed with covariance **Σ**, then ***a*** is also Gaussian distributed with mean ***a**_g_*. Under these assumptions, it follows that the sample mean is an optimal summary statistic:

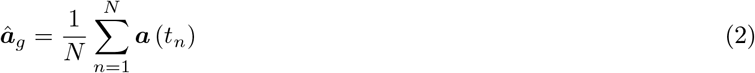

#### 2.2.2 Time-varying orientation

Assumption (2) above requires more sophistication as we cannot ignore time, which invokes the topic of *digital signal processing* [Little, 2019]. In particular, *digital filtering* methods attempt to solve the problem of recovering a signal “hidden” in the measurements, which we can write as a mathematical operation *H* [***x***] where ***x*** = (*x*_1_,…,*x_N_*) is a digital signal with *N* measurements sampled at equally-spaced time intervals. The simplest of such filtering methods are *linear-time invariant* (LTI) processing algorithms: that is, they satisfy the following two mathematical properties [Little, 2019]:

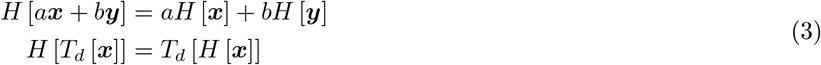

where *T_d_* indicates *time delay* by *d* time units and *a, b* are constants. Such methods can be entirely characterised using the *Fourier transform* (that is, by analyzing observed data in terms of frequency rather than time). In particular, assumption (2) suggests LTI *smoothing* or *low-pass filtering* of the measured signal to attempt to recover ***a**_g_*. To avoid mathematical complications, this should be carried out on uniformly-sampled signals of sampling interval Δ*t* seconds. Although it is possible to obtain highly regular sensor data from the Android smartphone operating system, this usually requires specialized (real-time) software engineering techniques to minimize the potential for large gaps between sensor measurements. Given sufficiently regular but non-uniform sampling, a simple approach (which we use) to convert irregularly sampled sensor data into uniformly sampled data, is *piecewise linear interpolation*; the measured data is replaced by:

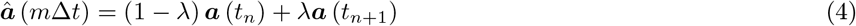

where *λ* ∈ [0, 1] and *m* = 1,…, *N*. For each *m*, the values of *λ* and *n* are chosen such that *m*Δ*t* ∈ [*t_n_, t_n+1_*].

Low-pass filtering involves, ideally, removing all frequency components above a certain frequency threshold. For example, one may want to completely remove fluctuations occurring at a higher rate (*cutoff frequency*) than, e.g. 0.5Hz (2 seconds duration). Unfortunately, this ideal is never achievable in practice due to (the Fourier equivalent of) Heisenberg uncertainty, so there is a trade-off between the accuracy with which the low-pass filter can remove fluctuations, and the effective resolution in time of the filtering. A very accurate low-pass filter will usually have unacceptably poor time resolution. The consequence is that time-localized features in the signal such as a stepping event, will become unacceptably “smeared out” in time after filtering.

Manipulating this trade-off leads to the discipline of *rational digital filter design* [Little, 2019]. Different digital filter designs manage this trade-off in different ways; a widely used filter for accelerometry processing is the low-pass *Butterworth filter*. This particular design has the useful property that it has constant amplitude response for signals below the cutoff frequency. However, as with all such *infinite impulse response* filters, different frequency components of the signal are shifted in time by different amounts, and this has a significant negative impact on the ability to properly localize events such as individual stepping events in the signal. Essentially, the fundamental problem with LTI digital filtering such as the low-pass Butterworth filter, and why we do not recommend it for practical applications in accelerometry processing, is that the signal we want to retain (smoothly varying orientation with occasional abrupt changes) and the events we want to remove (dynamic features such as step events occurring on a short time scale) overlap in the frequency domain, and there is no LTI filter which can separate them in principle [Little and Jones, 2011].

To get around these limitations we have to turn to some kind of *nonlinear* or *time-varying* filtering [Little, 2019] in order to separate the gravitational orientation signal from the interpolated accelerometry data. This is a very broad area of active research. *Discrete-time wavelet shrinkage* has found some use in this application, but Heisenberg uncertainty (here, the time-frequency trade-off is allocated differently to Fourier analysis), causes similar problems with stepping event localization and therefore we do not use it here. One of the simplest filters which is consistent with assumption (2) above and can, in principle, recover localized events and also perform gravitational smoothing, is *L1-trend filtering* [Kim et al., 2009]. This filter assumes that the gravitational component changes in a *piecewise linear* manner. This can be expressed as the following minimization problem:

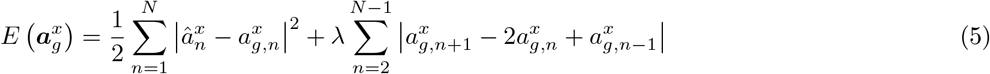

where *λ* > 0 is a regularization parameter which controls the timescale of the piecewise linear smoothing. Here, 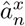 refers to the *n*-th interpolated, measured accelerometer observation in the *x*-coordinate; 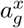 refers to the x-coordinate of the gravitational signal to be estimated, and 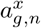, the *n*-th time element of that signal. There will be three such equations (5), one each per *x, y, z* coordinate.

To solve the above problem, we minimize 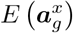 with respect to 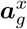 (and similarly, for each of the other two coordinates, *y, z*). The problem is globally *convex*, thus, there is a unique solution which can be found using a variety of numerical optimization methods [Little, 2019]. The first term in this expression is the sum of squared errors representing the “fit” of the estimated piecewise linear gravitational signal 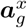 to the measured signal 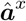. The second is the sum of the magnitude of (discrete) second derivatives in time. Noting that 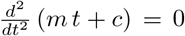, we can see that the second term encourages a solution which has minimal total magnitude of changes in gradient. This means that, approximately, the solutions to (5) are piecewise linear.

#### 2.2.3 Body-frame coordinates

Having estimated the gravitational orientation vector (in all three dimensions), the dynamic acceleration due to the user’s movement can be estimated simply by computing 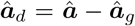. This will suffice for many applications where we do not need to know anything more about the physical meaning of the resulting vector; for example, as we discuss below, we use this simple procedure for step rate estimation to inform the choice of prior distribution.

In other applications where more detail about the dynamic acceleration of the user’s body is required, it is necessary to somehow give an interpretation of the measured coordinate frame relative to the body. This is the case for detecting individual stepping events. A frame which is highly relevant for walking tests has the dynamic *z* coordinate aligned with the longitudinal axis (up-down) since we expect the user will be standing upright with gravitational force acting ‘downwards’, the *x* coordinate aligned with the anterioposterior axis (front-back) since we anticipate the user will be walking in a straight line, and the *y* coordinate aligned with the frontal axis (left-right). Clearly, other coordinate permutations are possible and entirely equivalent and indeed this is an inevitable ambiguity we must resolve.

Given 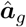, an appropriate set of three-dimensional rotations can perform this centering. To do this, we first need to convert the Cartesian coordinates into *spherical* coordinates, *angle g_ϕ_* and *elevation g_θ_*:

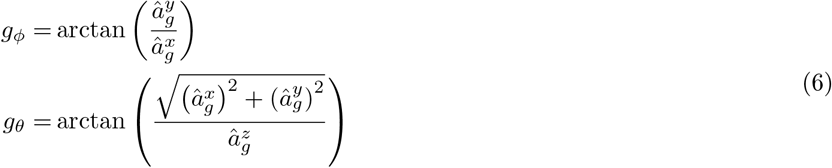

from which we can write down the corresponding *rotation matrices* around the *x* and *y* axes, transforming the origin to this estimated angle and elevation:

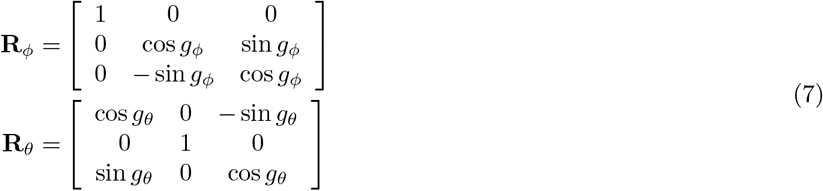

We can then undo the effect of these rotations, to find the dynamic acceleration in the body-centred frame, 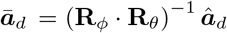. An important consideration is that the above coordinate transformation does not distinguish the *sense* of the placement of the IMU. Indeed, the measured coordinates can be permuted and/or reversed in sign and yet (6) computes only the *principal value* of the spherical coordinates. As an example, reversal of the sign of 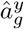 cannot be detected if 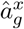 is simultaneously reversed. Knowledge about the sense of the measurements is required to resolve this permutation ambiguity.

One approach to sense correction is to establish the context of each measured coordinate by integrating over time. There are many ways to do this; the method we have developed relies on the determination of the minimum and maximum values of the rotated *x* and *z* coordinates during the walking test. Due to the nature of the walking test protocol, one coordinate incorporates the acceleration due to gravitational reaction force which is in practice, readily separated from the other coordinates by the predominant value ranges. This allows sense correction by appropriate coordinate permutations and/or sign reversals. More sophisticated methods may be necessary if the users can deviate more substantially from the test protocol.

### 2.3 Discriminating walk from rest

In the context of a walking test, it is important to be able to detect whether an individual is walking or is at rest. If at rest, then clearly any steps “detected” will be spurious. Provided the individual follows the particular 6 minute walk test protocol, then we only expect two behaviours: walking and at rest. These will then be distinguishable from the accelerometer. In particular, if the individual is at rest, then the dynamic acceleration magnitude will be small. By contrast, during walking we expect significant changes in kinetic energy which should be detectable in significant magnitude of acceleration. These simple criteria suggest that clustering the dynamic acceleration magnitude into two states (walk versus rest) should suffice.

Treating the dynamic acceleration magnitude as a random variable, the two-state model is, formally, a *mixture distribution* [Little, 2019]. Mixture distribution estimation is a large topic covering techniques such as Gaussian mixtures fitted using expectation-maximization (E-M) and hidden Markov modeling (HMMs). This level of complexity is not particularly well justified in this setting, therefore, we take a simpler approach to estimating this pair of distributions.

The pre-processed (interpolated, range normalized, trend filtered) dynamic acceleration signal is split into windows of 3 seconds, overlapping by 2.7 seconds (these window durations are chosen for a good balance between robust magnitude estimation and sufficiently rapid response to changing behaviour in this walking test; for other applications we might expect a different trade off). For each window, the standard deviation of the dynamic magnitude is computed, and used to represent the (window-averaged) dynamic magnitude 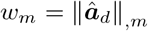 where 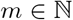 is the window index. We next assume that both walk and rest distributions are Gaussian with pre-specified standard deviation *σ* = 0.2 (chosen to represent typical variation in dynamic magnitude). Assuming that the window with least motion recorded corresponds to rest, we set the most likely rest mean to be 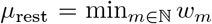. Also, assuming that the typical level of motion corresponds to walking, we choose 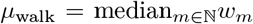. Implicit in these assumptions is that the minimum level of motion in each window is substantially different from the typical level, this is generally true if adherence to the walking test protocol is good. The explicit model for each of the two distributions is:

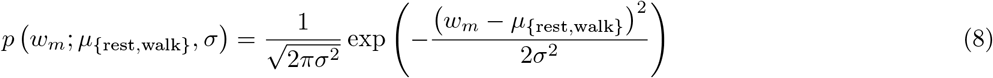

Given these two models, it is straightforward to “decode” the state *s_m_* ∈ {rest, walk} of each window m by computing the probability density of *w_m_* under both models and maximizing:

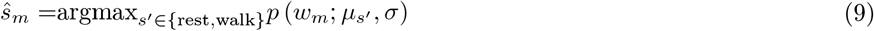

which can be shown to be equivalent to minimizing the square of the difference between the mean for rest and walking states [Little, 2019], 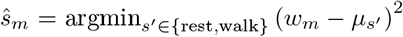. The Bayesian posterior probability of each window state will be useful later; it is obtained using the following computation [Little, 2019]:

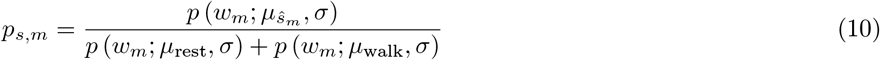

### 2.4 Detecting discrete step events

Walking is a cyclic activity which can be decomposed into “phases”: the stance phase and the swing phase on each side, themselves composed of discrete events such as the heel strike and toe-off events. Under controlled lab conditions, it is possible to identify these separate phases and events and quantify their properties. Outside the lab, however, the variability of numerous uncontrollable factors such as device placement, robust physical contact of the device with the body, walking surface, footwear and obstructions on the walking route, and the interaction between intentional and unintentional behaviours, mean that detailed quantification of these phases and events is often very difficult.

In the body-centred coordinate frame described above, accurate lab-based measurements allow the quantification of the dynamic force and/or acceleration trajectories during the typical gait cycle [Vaverka et al., 2015]. From these measurements it is easy to identify each step. The goal of algorithm development in this context is to extend this to wearable IMU measurements in uncontrolled conditions, a much more difficult task. At the least, however, provided adherence to the walking protocol is sufficiently good, it should be possible to uniquely identify the cycle on each side (left/right), which contains an individual step per side. We refer to this as *step event* detection.

To address this, we use a simple *morphological matching* method which attempts to locate a canonical template model of the acceleration profile during each gait cycle. The model has the analytical form:

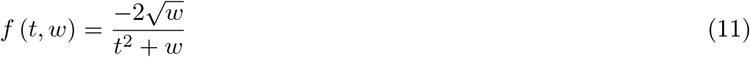

with width *w* > 0. The maxima and minima are located at 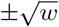 where the function takes on a peak value of ±1, and the function goes through zero at t = 0. As w increases, the swing through zero becomes faster. To a close approximation, this is the shape of the forward/backward acceleration during each side’s gait cycle. The positive peak corresponds to the propulsive (*push-off*) phase of the cycle when forward momentum is imparted, and the negative peak the braking phase [Vaverka et al., 2015].

To match the template to the signal, we use the *cross-correlation* method. This can be understood as “sliding correlation”, that is, the correlation between the (short-time) template and the signal is computed at all time delays [Little, 2019]. First, the template is sampled at the same uniform rate Δ*t* as the pre-processed, body-frame centered forward/backward acceleration signal 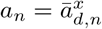 to obtain the template signal:

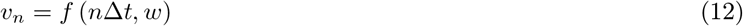

for *n* = –*N*_2_,…, *N*_2_ where 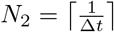. Then, these two signals are cross-correlated:

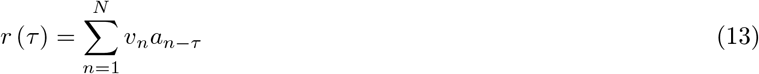

for *τ* = –*N*_2_,…, *N*_2_. Any terms in the cross correlation sum where *n* – *τ* is outside the range of the pre-processed accelerometry signal, are set to zero.

Next, a decision needs to be made about the location of each step event. For this, each local maximum in *r* (*τ*), *τ*_max_ = arg max*_τ′_ r* (*τ′*), must be assessed for its likelihood of representing a possible “match”. There are many ways to do this using techniques of varying levels of sophistication, but in experiments, the following very simple rule was determined as most effective across all the data collected for this protocol:

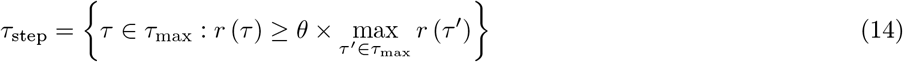

where *θ* = 0.7. In words: step event times *τ*_step_, are selected from the local maxima for the cross-correlation, where the cross-correlation at these local maxima are sufficiently large, relative to the maximum cross-correlation. Given the individual step event times *τ*_step_, it is straightforward to derive cumulative properties such as the number of steps and the median (typical) time interval between steps, and the range of intervals.

These estimated step events will also play a crucial role in Bayesian ‘anchoring’ of step rate estimates, discussed next.

### 2.5 Bayesian step rates

Step events implicitly decompose walking activity into discrete, component events. However, walking is, in reality, rarely an *intentionally* composite ‘stepping’ activity. Rather, it is better to understand normal walking as a continuous, instinctual activity, where the instantaneous *rate* of cycling through the sequence of muscle activations is modulated according to intentions, in service of larger goals. From this perspective, the individual events are not important, and the aim is to quantify the rate of walking from the accelerometry signal. For example, in many movement disorders, the individual events in the gait cycle can become very distorted and idiosyncratic, but the activity is still clearly repetitive.

This problem is best addressed using some kind of *autocorrelation* method which can be understood as *morphological self matching* at all time delays. Traditionally, autocorrelation is just a special case of cross-correlation (13), but there are some practical weaknesses which suggest the use of slightly different measures of similarity. In particular, one issue with autocorrelation is that it varies *smoothly* with changes in time delay (corresponding to step rate). This may seem mathematically convenient but this makes it difficult to uniquely identify the most probable time delay because traditional autocorrelation spreads information about the optimal self-match over many time delays. For our purposes, a more useful measure of self matching for uniformly sampled time series is the *mean absolute self mismatch*:

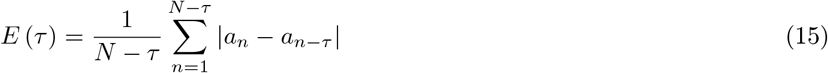

where 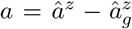, the *z* coordinate of the estimated dynamic acceleration after trend filtering (5). This measure has similar behaviour to autocorrelation, but with the following properties:

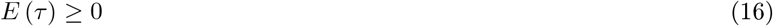

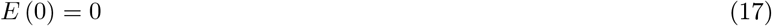

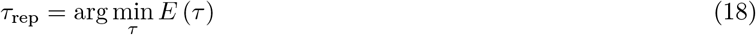

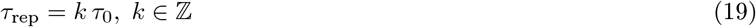

The last condition states that all minima *τ*_rep_ of repetitive signals a, are integer multiples of the *fundamental period τ*_0_. More importantly, the measure does *not* smoothly vary with *τ* so that the minima of *E* (*τ*) are highly localized and thus easy to isolate. By contrast, maxima of the autocorrelation are inherently ambiguous. In this application, because we need high-precision estimates of each value of *τ*_rep_, we use a local quadratic polynomial model for each minima using three local time delays (i.e. based on the *E* (*τ*_rep_ – 1), *E* (*τ*_rep_), *E* (*τ*_rep_ + 1)), fitted using least-squares. From this quadratic polynomial *E* (*τ*) ≈ *aτ*^2^ + *bτ* + *c* with parameters *a, b, c*, the predicted location of the minima is just 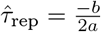.

To identify step rates, we first note that (15) really only makes sense as a *stationary* measure, that is, it only has a straightforward interpretation under the assumption that the gait behaviour is (almost exactly) repetitive over time. This is unrealistic as in practice gait behaviour patterns are always constantly changing. To partially address this problem we use the windowing described in Section 2.3, so that the computation of step rates is obtained for each window making the analysis *piecewise* stationary.

Next, to identify step rates, a naive choice would be the smallest minima of (15) (excluding *τ* = 0). However, this criteria has an inherent ambiguity because condition (19) indicates that the global minima for E may coincide with *E* (*kτ*_0_) for some *k* > 1. For example, for gait signals in this application, it is often the case that the signal is highly symmetric with respect to left/right sides. In this situation, the smallest 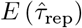 represents the partial gait pattern involving each side separately. Whereas, if the signal is asymmetric (because the device is, say, placed on one hip rather than at the small of the back), then the smallest 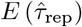 represents the full duration of the gait cycle of both sides. This ambiguity is a significant unsolved problem with periodicity analysis in general, arising in, for example, speech and electrocardiogram signal processing [Shi et al., 2019, Sun, 2000].

The only way to resolve this ambiguity is to bring in other information, and there are many possibilities here. A very common resolution is to assume that the gait cycle can only assume a certain range of durations. This is not unreasonable – human gait is constrained by physiology so that a gait cycle of 0.05 or 5.0 second’s duration is obviously impossible, for example – yet cycle durations of 0.6 (*k* = 1) and 1.2 (*k* = 2) seconds are entirely plausible and so cannot practically be separated in this way. Thus, we clearly need a more sophisticated algorithm. In order to achieve this, we exploit information from signal morphology, in particular, the step events detected in the previous section above. The timing of detected step events allows a rough prediction of the stepping rate, and can be used as a *probabilistic constraint* on the step rate estimate to resolve the periodicity multiple ambiguity.

A mathematically consistent and principled approach to applying such constraints, is the use of *Bayes’ rule* [Little, 2019]:

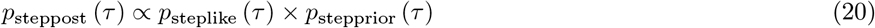

This is a formal statement that applying the constraint (*prior*) to the measured value (*likelihood*) creates a new distribution *p*_steppost_ (*τ*) (*posterior*) which is proportional to the product of the distribution over the constraint *p*_stepprior_ (*τ*), and the measured value *p*_steplike_ (*τ*).

#### 2.5.1 Step rate prior

Given information from the step events detected above, we can construct a prior distribution over the step rates. This distribution is a matter of design choice, and for simplicity we use an *exponential power* model which represents a useful trade-off between narrowing down the step rates to the step event estimates, and allowing sufficient flexibility since the step event estimates are themselves not perfect:

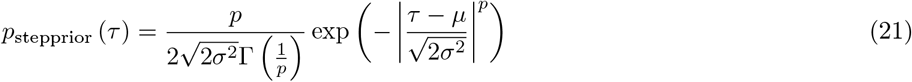

where *p* > 0 is a “sharpness” parameter which determines how narrow the distribution is around the most likely value 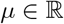. The parameter *α* > 0 acts much like a variance (i.e. spread) parameter, and when *p* = 2, this distribution coincides with the Gaussian (normal) distribution which has a fairly narrow peak around *μ*. To spread out this peak, thus relaxing the constraint, we set *p* = 10. We centre the peak of the prior around the typical step event rate, *μ* = 2×median (*τ*_step_). Finally, we choose the spread parameter to be proportional to the uncertainty in the step event rate estimates, *σ* = 2 × iqr (*τ*_step_) where iqr is the *interquartile range*, which is the difference between the 75th and the 25th percentile.

It should be pointed out that there are cases where the step event detection fails, particularly when the gait test protocol is significantly violated. This can be detected by analysis of the step event data. For example, the step events may be extremely irregular, physically implausible, or there may be no detected steps at all. In these cases, it may be much more reliable to fix the parameters *μ* and *σ* in the prior (21).

#### 2.5.2 Step rate likelihood

Having selected a step rate prior distribution, we need a distribution for the likelihood *p*_steplike_ (*τ*) in order to compute the posterior. This requires finding a model for the distribution of 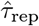 in such a way as to extract and summarize the relevant details about the periodicity in the signal. Because this distribution is expected to be *multimodal* (it has multiple most likely values), it would be very difficult to find a simple and useful parametric model for the distribution. Therefore, we have little option but to resort to a non-parametric *kernel density* distribution [Little, 2019], here, using a Gaussian (normal) kernel for simplicity. Such nonparametric distributions typically require quite large amounts of data in order to get reliable estimates, particularly for rare events. In this situation we only have (on the order of) 100’s of step rate estimates, *τ*_rep_ which may leave some of the rarer step events with unreliable distribution estimates. One solution is to use the mismatch *E* (*τ*_rep_) as a measure of the importance to attach to each step rate estimate. This has a natural interpretation as the *weight* of each of the *N_τ_* step rate estimates in the kernel density:

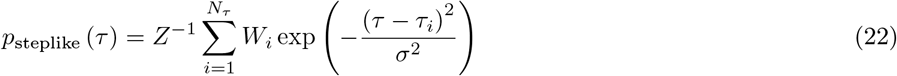

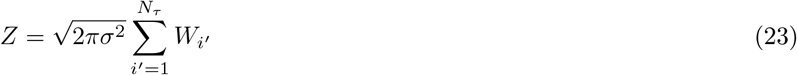

To convert the measure of mismatch *E* (*τ*_rep_) into a weight *W* which is increasing with increasing importance, we rescale the mismatch by maximum mismatch:

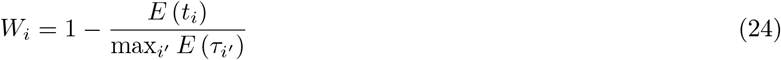

which leads to weights in the range 0 ≤ *W_i_* ≤ 1.

#### 2.5.3 Derived posterior

The final step in estimating step rates is to compute the posterior distribution, *p*_steppost_ (*τ*). Since the likelihood and prior distributions are mixed analytical (parametric) and non-parametric so that there is no simple analytical formula for the posterior, we evaluate the posterior numerically. This is done by placing a fine grid of *N_Δτ_* = 1, 000 intervals over the maximum range of the step rate, here set to biologically plausible values of 0 ≤ *τ* ≤ 3.0 seconds. This choice of evaluation grid parameters leads to a step rate resolution of 0.003 seconds. Of course, a finer grid would give better resolution which might be necessary for some applications. The most (posterior) probable (*maximum a-posteriori*, MAP) value of the step rate, which is important for practical purposes, can then be found by maximizing the posterior over the evaluation grid:

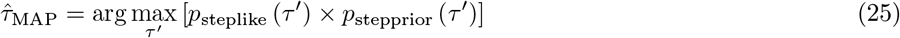

This MAP value, along with the corresponding probability (density) value 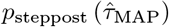, are useful in practice as they give a simple summary of the posterior rate and its associated probability. However, this ignores the uncertainty in the rate estimate, which is critical for further statistical analysis. For instance, the posterior could be sharply peaked around the MAP value, or it could be at the top of a shallow hill. The former would inspire high confidence in the estimate, but the latter would imply that the estimate is of low reliability. There are many ways in which this uncertainty can be quantified, for example, a *confidence interval* gives a range of values into which 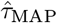 falls, which can be computed from the *cumulative* posterior distribution, 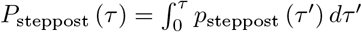.

### 2.6 Estimating distances and speeds

Whilst IMU devices such as accelerometers and gyroscopes are well-suited to estimating detailed information about step quality and quantity, it is much more difficult to estimate larger-scale features such as walking speeds. This is because IMUs are *differential* devices designed to quantify spatially (and temporally) localised information about inertial frames. To get from local to more global inertial information, it is necessary to perform some kind of *integration* over long time scales, but errors in locally-estimated quantities such as speed accumulate, such that they come to dominate the quantities of interest. For this reason, a much more reliable solution is to rely on *global* localisation such as GPS devices embedded in modern smartphones.

### 2.7 Pre-processing GPS data

GPS receivers are based on the principle of *distance triangulation* from orbiting satellites which transmit wireless signals received by the GPS device in the smartphone. The position in orbit of each satellite is known. Therefore, the distance from the GPS receiver to each satellite can be inferred from the time of arrival of the wireless signal, knowing the speed of light. This timing information, which must be highly precise to produce sufficiently accurate distance measurements, is provided by atomic clocks located on each satellite and an internal clock in the receiver. Using at least three distances, it is possible to localize any point on the Earth’s surface, but the lack of precision of the GPS receiver’s clock means that a fourth measurement is normally required to eliminate the GPS receiver’s time offset from the distance estimation computations. Since each satellite provides one distance measurement, the more satellites available the better the distance measurements. The US GPS system, upon which a substantial fraction of the world population relies for location tracking, has around 30 or so working satellites in an orbital configuration such that up to 6 satellites at a time can, in principle, be received from anywhere on the surface of the Earth. In theory then, this setup provides ample, high-precision triangulation measurements to estimate latitude, longitude and elevation to within a few metres, continuously.

Unfortunately, GPS wireless signal reception is far from ideal in reality. The wireless signals are not very powerful and can only be reliably received along a direct, line-of-sight path to the satellite. This generally rules out GPS location estimation indoors, for instance. It also means that very common sources of interference include urban structures such as high-rise buildings in built up areas. Much more insidious than direct path signal degradation are “phantom” paths induced by objects which cause secondary reflections and refraction, which may masquerade as the direct path, artificially lengthening the estimated distance to the satellite. In practice then, GPS location tracking does not use the “raw” GPS position measurements, because these are generally patchy with multiple missing and/or inaccurate measurements. Instead, these need to be pre-processed and interpreted in such a way as to make them useful for practical location tracking applications.

GPS units generally report a position in standard global spherical co-ordinates such as longitude and latitude e.g. 51.77508844634028° north, −1.279890732938649° west, 105 metres above sea level. For our purposes, where we need to measure distances on the order of a few hundred metres, then a more local Cartesian co-ordinate system such as UTM (Universal Transverse Mercator), constructed of multiple overlapping, *conformal* (angle-preserving) maps, is more useful. The mapping from global co-ordinates to the local UTM map we use (WGS84 spatial reference system) is based on an analytical model of the Earth’s surface (oblate spheroid) and is accurate to a few millimetres [Karney, 2011].

### 2.8 Walking test distance estimation

After conversion to UTM horizontal and vertical co-ordinates (metres) which we denote by 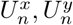 at each location measurement’s time instant 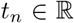, the next step is to estimate the *path length* of the GPS measurements, since we are not interested in the absolute location. Probably the simplest approach to doing this is to calculate the displacement vector between each measurement:

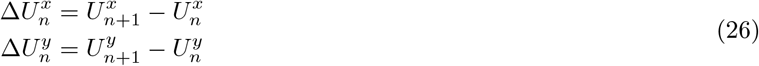

from which the path length *d* can be easily estimated as the sum of the lengths of each of these vectors:

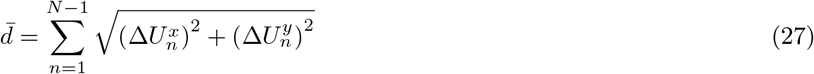

The main assumption behind this simple distance estimate is that, in between GPS location measurements, the walking path of the test subject is a nearly straight line. This is fairly realistic approximation given the test protocol, provided that the typical time interval between measurements, Δ*t_n_* = *t_n+1_* – *t_n_*, is sufficiently small (on the order of 1 second), and the location measurements are accurate (to within a metre or so). Experimentally, we have found that, provided the GPS unit reports good quality signal (that is, it has line-of-sight access to a handful of satellites), then these assumptions are satisfied in the main. In highly built-up areas, these assumptions may not be true and we may need more sophisticated distance estimation algorithms.

### 2.9 Instantaneous speed estimation

Although GPS units do have internal “tracking” algorithms, using which they can be made to report speed estimates, these algorithms are not sophisticated enough to be scientifically reliable. Instead, it is necessary to process the location data in order to estimate (instantaneous) speed. Velocity is position differentiated by time, so speed, being the magnitude of velocity, requires estimating this differential as accurately as possible from the (somewhat unreliable) GPS location estimates.

Numerical differentiation is an extensive topic, see for instance Press et al. [2002], but the essential idea is that the differential of position, *dU/dt*, is a numerical difference, for example:

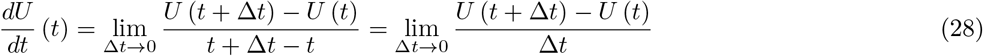

which converges on the true velocity in the limit as Δ*t* goes to zero. The main problem is that we do not have an analytical form for the position, *U* (*t*), so that the limiting differential is not computable in this setting. Instead, we have to resort to approximating this differential using some kind of difference. A simple approach is *Euler’s* (*forward*) *method*, which assumes that the above difference is a close approximation to the differential:

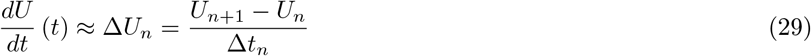

where Δ*t_n_* is the time difference between location measurements.

If the changes in distance are small by comparison to the size of the location errors, and the time differences are small, then this estimate will inevitably exaggerate the location errors. Combined with satellite drop-out, this means it is critical to integrate this estimate over a longer period of time. This again raises the familiar problem of balancing responsiveness of the estimate (it should reflect local changes in position) against the need to identify and correct for GPS location errors. The classical approach to addressing this dilemma is to use some kind of *tracking* algorithm, for example, the (linear, Gaussian) *Kalman filter* [Little, 2019] which is based around a forward, linear model of predicted device movements. It computes an optimal, Bayesian balance of the uncertainty in the predictions against the uncertainty in the GPS measurements as they become available. There are weaknesses to this algorithm, in particular, its tendency to get stuck in overshoot/undershoot cycling when the device changes direction rapidly, for example. This is why, in road vehicle navigation applications, Kalman filter tracking is generally projected onto a known map.

Although we do not have access to cartographic constraints because the walking test could be conducted anywhere outside, in our case, we can exploit the fact that we have the full record of GPS observations available at the time of processing the gait test data (the *offline* situation). This means that we can integrate both forward and backward in time to smooth out location errors while maintaining responsiveness to rapid changes in position. A mathematically principled solution to this offline smoothing problem is *Gaussian process* (*GP*) *regression*, which can be conceptualized as a highly generalized kind of Kalman filtering whereby information about the changes in location across the entirety of the GPS record, can be assimilated in a way which is consistent with Bayes’ theorem [Little, 2019]. The trade-off between smoothing and responsiveness is determined by a choice of prior *covariance function *κ*_0_* (*t, t′*).

Given the speed estimated from the time differenced GPS location data (29) *u_n_* = ∥Δ*U_n_*∥ and based on a squared exponential prior covariance function, the mean and (co)-variance of the speed can be computed at any time instant 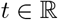:

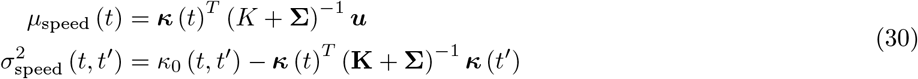

where the *kernel matrix* has entries *K_n, n_′* = *κ*_0_ (*t_n_, t_n_′*), the *kernel vector κ_n_* (*t*) = *κ_0_* (*t, t_n_*) and the likelihood covariance is a diagonal matrix whose values Σ*_n, n_* are set equal to the reported GPS location uncertainty. Such GP regression amounts to finding a Gaussian distribution for each time instant, so the distribution of speed *u* (*t*) at time *t*, is estimated as:

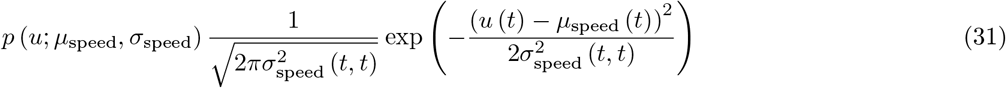

From this, it is straightforward to construct a confidence interval for the speed estimate at any time instant.

Experimentally, we have found that for smartphone GPS units, the shortest time differences between location measurements is about 1 second. Therefore, short resting periods during the walk test may not be readily represented in the speed estimates above. For this reason, when computing the overall average speed for statistical analysis purposes, any rest periods are removed first.

## 3 Smartphone application development

We used a prototype app containing a proprietary sensor data collection module developed by Signant Health (Blue Bell, PA, USA) to collect accelerometer and GPS data to be used in algorithm development. This module was designed to detect all sensors on-board an Android smartphone, and exposed these via an interface that enabled a performance outcome test developer to select which sensors to sample from, the sampling rate for each, and the duration of the sampling period (Figure 1). The app ensured that the on-board GPS was turned on when the app was opened to ensure that adequate satellite lock was achieved when the test commenced. The app provided an audible countdown timer to mark the start of the test, and an audible tone on test completion. At the end of each test, data were compressed and securely transmitted to a cloud-based data repository. This sensor data collection module was subsequently incorporated into a patient-facing app that is intended to be used to administer a short (six-minute) walking performance outcome test in patients at home (Figure 2).

**Figure 1:**
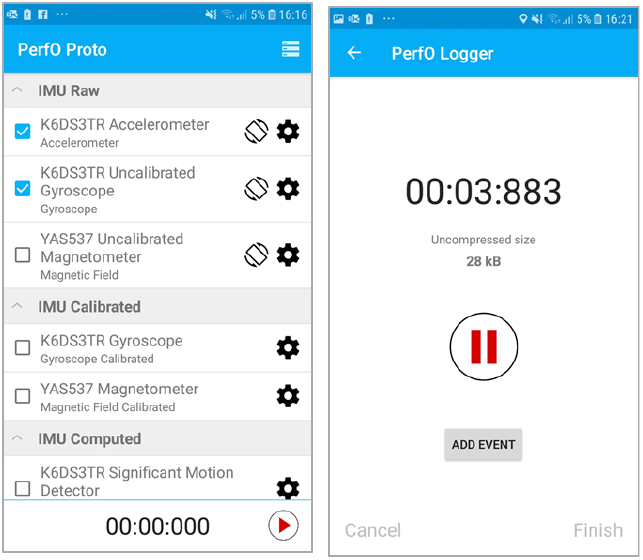
Smartphone application interface to set up sampling from any on-board sensor for performance outcome test data collection. (Left) Interface for sensor selection, (right) countdown screen during walking test conduct.

**Figure 2:**
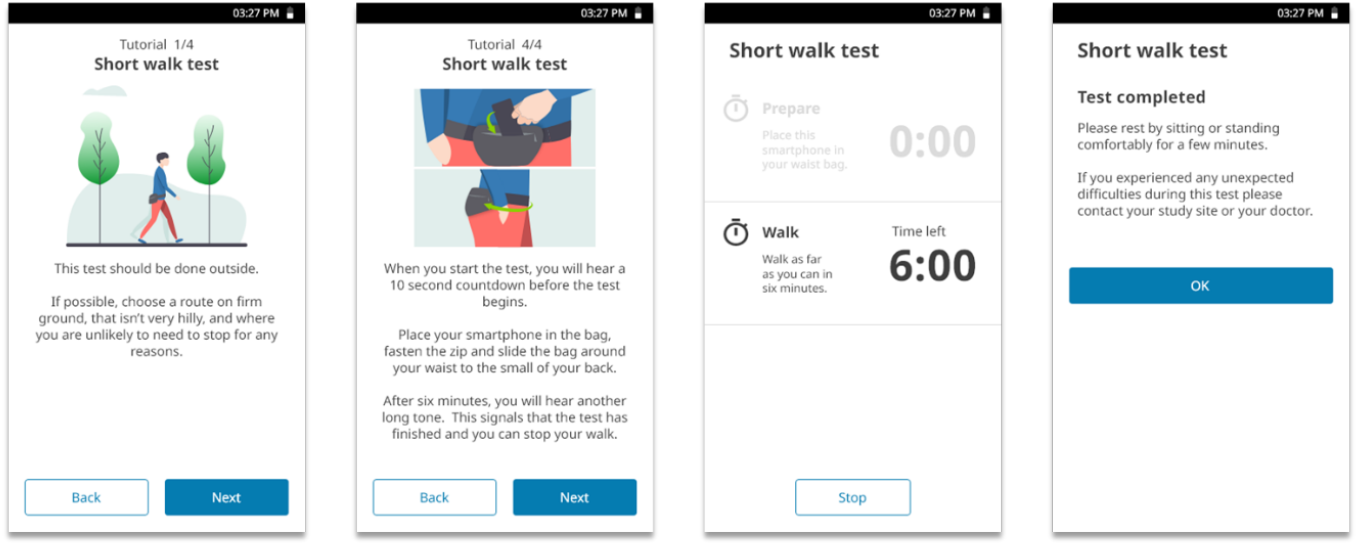
User interface for the smartphone short walk test.

## 4 Data collection

Data for algorithm development were self-collected by the authors, all members of the Signant Health clinical outcome assessments science team. During each test, actual distance covered and number of steps completed were recorded by an independent observer. Actual distance covered was measured to the nearest cm using a Trumeter 5000 Professional Road Measurer (Trumeter, Coconut Creek, FL, USA). The number of steps completed was estimated by manual counting using a manual tally counter that was clicked once per observed step by the observer.

The app was installed on a Samsung Galaxy A20 smartphone (Samsung, Seoul, South Korea). This was placed in landscape orientation in a zipped waist bag that was worn to enable the smartphone to be located over the small of the back. Data were sampled from the smartphone accelerometer and GPS at rates of 50 Hz and 1 Hz, respectively.

Data under a number of different experimental conditions were collected, as outlined in Table 1. For subsequent analysis reported in the results section below, experiments #1 through #6 were merged, to obtain a total of 59 accelerometer time series from which to estimate step properties (one trace was corrupted and therefore unusable). Only GPS data from experiment #7 was used to estimate speed and distance computations. Experiments #1 to #6 used an early version of the app which activated GPS at the start of the walking test, which led to collection of a period of data prior to achieving adequate satellite lock. A further revision of the app (used in experiment #7) ensured that satellite lock was achieved before the walking test commenced.

**Table 1:** Summary of experimental data collected for the purposes of algorithm development. All experiments had no rests (continuous walking) except experiment #3 having two rests of 10 seconds duration at 2 and 4 minutes. All experiments were conducted in London, UK except experiment #7 conducted in Helsinki, Finland. Experiments #1 and #7 had two walkers, and experiment #5 had 4 walkers, 2 each of different age ranges (18-30, 31-65), the remaining experiments had only 1 walker. All experiments located the phone in a waist bag over the small of the back, except experiment #4 where additionally hand, shoulder and thigh pocket locations were used.

| Experiment # | Duration (min) | Surface | Pace | Repeats |
| --- | --- | --- | --- | --- |
| 1 | 2 | Flat: hard | Slow, medium, fast | 12 (6 per walker, 4 per pace) |
| 2 | 2 | Flat: hard, grit, grass | Slow, medium, fast | 18 (6 per pace, 6 per surface type) |
| 3 | 6 | Flat: hard | Slow, medium, fast | 9 (3 per pace) |
| 4 | 2 | Flat: hard | Average | 12 (3 per phone location) |
| 5 | 6 | Flat: hard | Average | 12 (3 per walker) |
| 6 | 6 | Hilly | Slow, medium, fast | 3 (1 per pace) |
| 7 | 6 | Flat: hard | Average | 16 |

## 5 Results

Estimation errors using the algorithms described above, were computed according to the following metrics: *mean absolute* (MAE) and *root mean square error* (RMSE) and their *percentage-normalized* (e.g. MAPE, RMSPE) and *posterior weighted* (e.g. WMAPE, WRMSE) variants. Specifically, for *N* predicted values *v_n_, n* = 1,2,…, *N* (e.g. steps per minute, speed, distance) and associated posterior probabilities *p_n_*, the total estimation error is calculated according to the general formula:

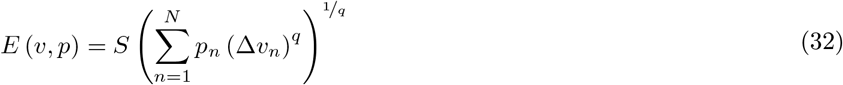

where, the positive estimation difference for each individual estimate is defined as 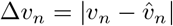 (scale factor *S* =1) and 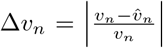 (scale factor *S* = 100) for relative (percentage) errors. For the absolute metric (MAE, MAPE, WMAE, WMAPE) q =1 and for root mean square metric (RMSE, RMSPE, WRMSE, WRMSPE) we set *q* = 2. Non-weighted errors (e.g. MAE, RMSE) set *p_n_* = *1/N*.

### 5.1 Signal processing

For accelerometry data, we select two sample time series to illustrate the function of the signal processing algorithms described above. Figures 3 and 4 show the effect of body-frame (7) and gravitational removal (5) transformations. These plots also show the locations of the step events estimated using (14). The corresponding continuous MAP stepping durations 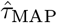, see (25) and posterior probabilities 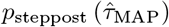 using step events to inform the choice of prior, are displayed in Figures 5 and 6. For the first example time series, we also show the distribution of the step event rate *τ*_step_, see (14), from which the prior model parameters are estimated (Figure 7). Finally, for the second example we show the estimation of walk/rest state from the dynamic acceleration amplitude (Figure 8).

**Figure 3:**
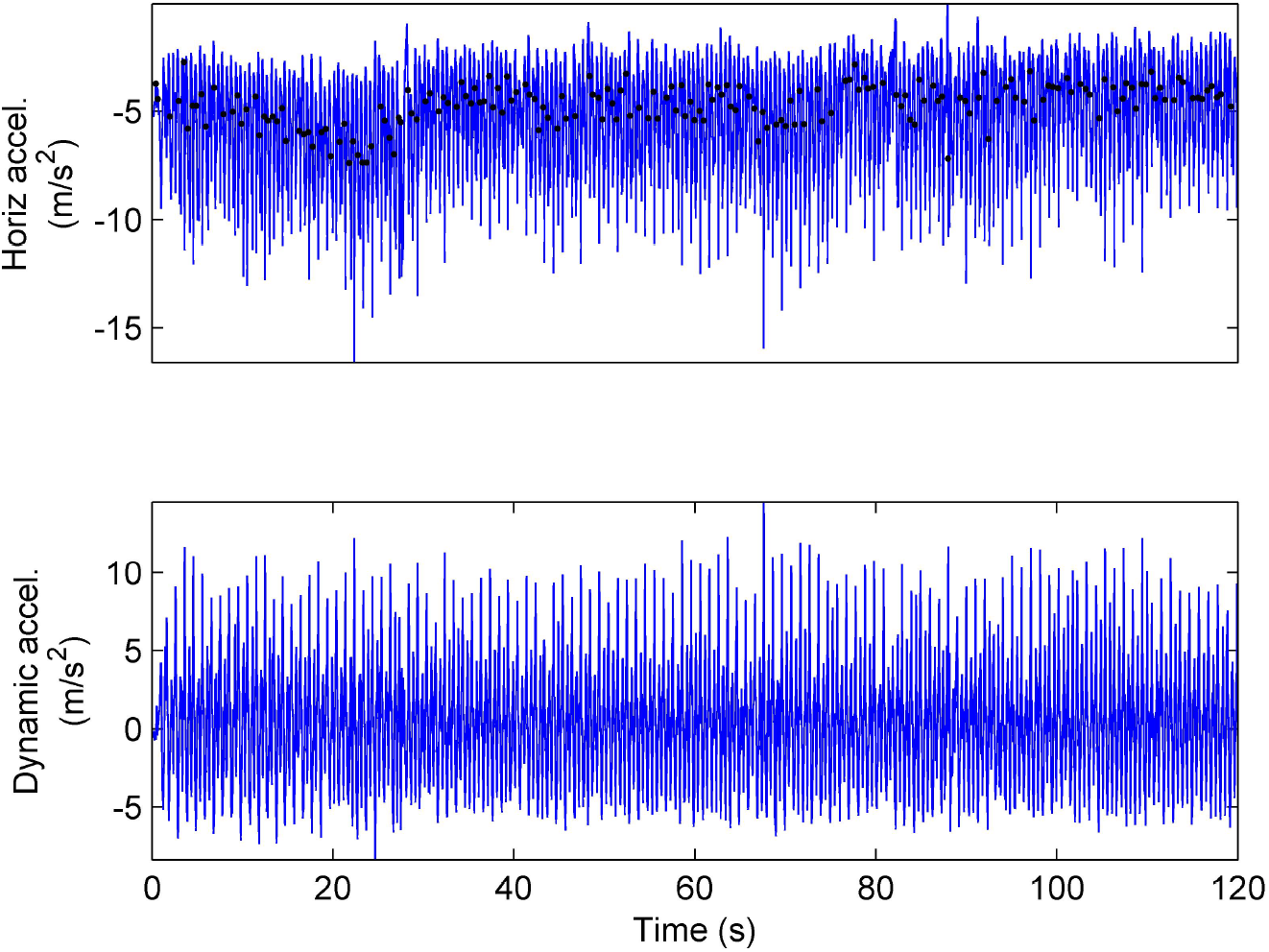
Example accelerometer data (blue lines) from experiment #4 processed to obtain the horizontal body-frame coordinate (top), along with the location of estimated step events (black dots), and the dynamic acceleration after removal of estimated gravitational component (bottom). Average pace on a hard, flat surface, with phone located in the hand of the wearer.

**Figure 4:**
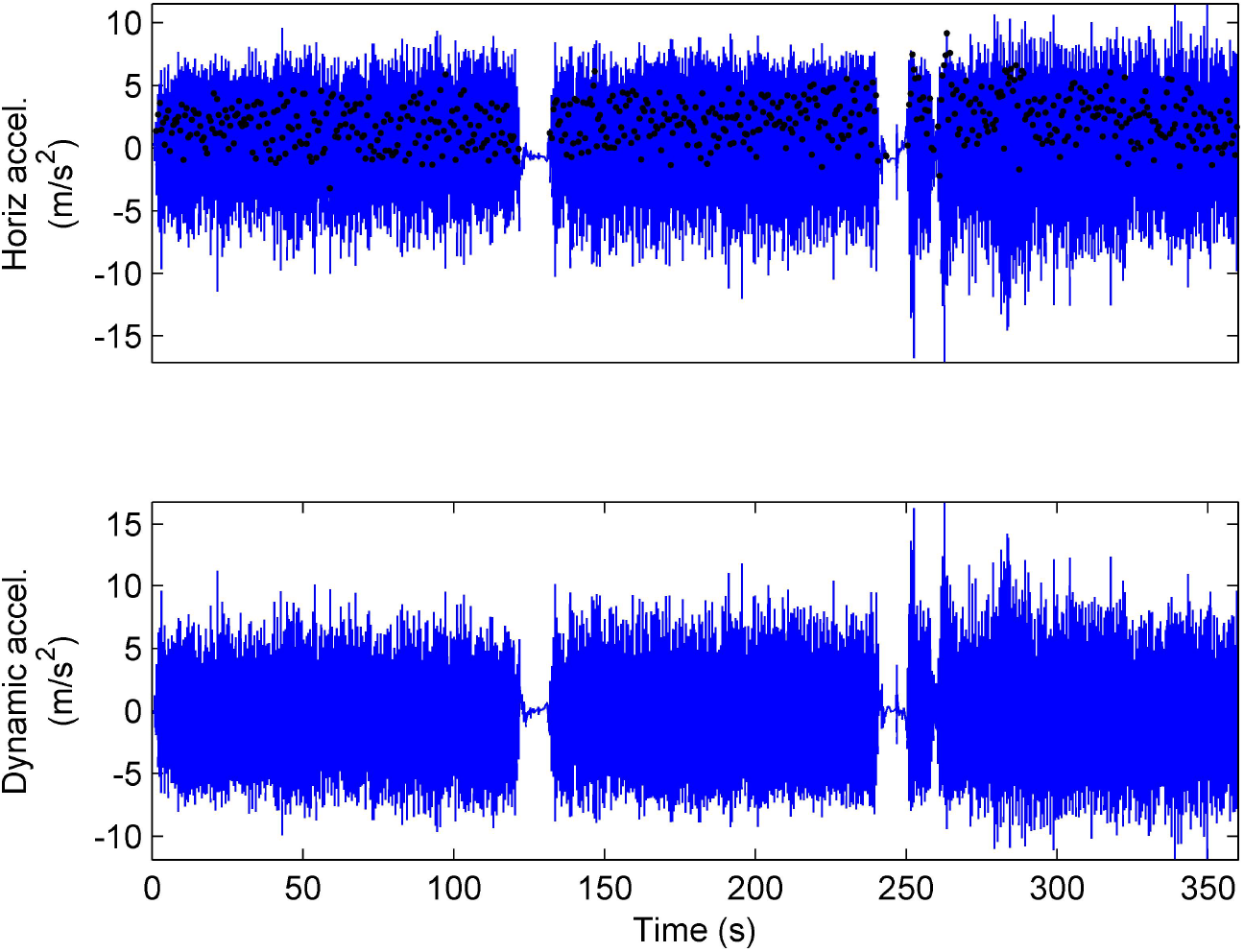
Example accelerometer data (blue lines) from experiment #3 with rests, processed to obtain horizontal body-frame coordinate (top), along with estimated step events (black dots), and the dynamic acceleration after removal of estimated gravitational component (bottom). Normal pace on a hard, flat surface, with phone worn in waist pouch at the small of the back.

**Figure 5:**
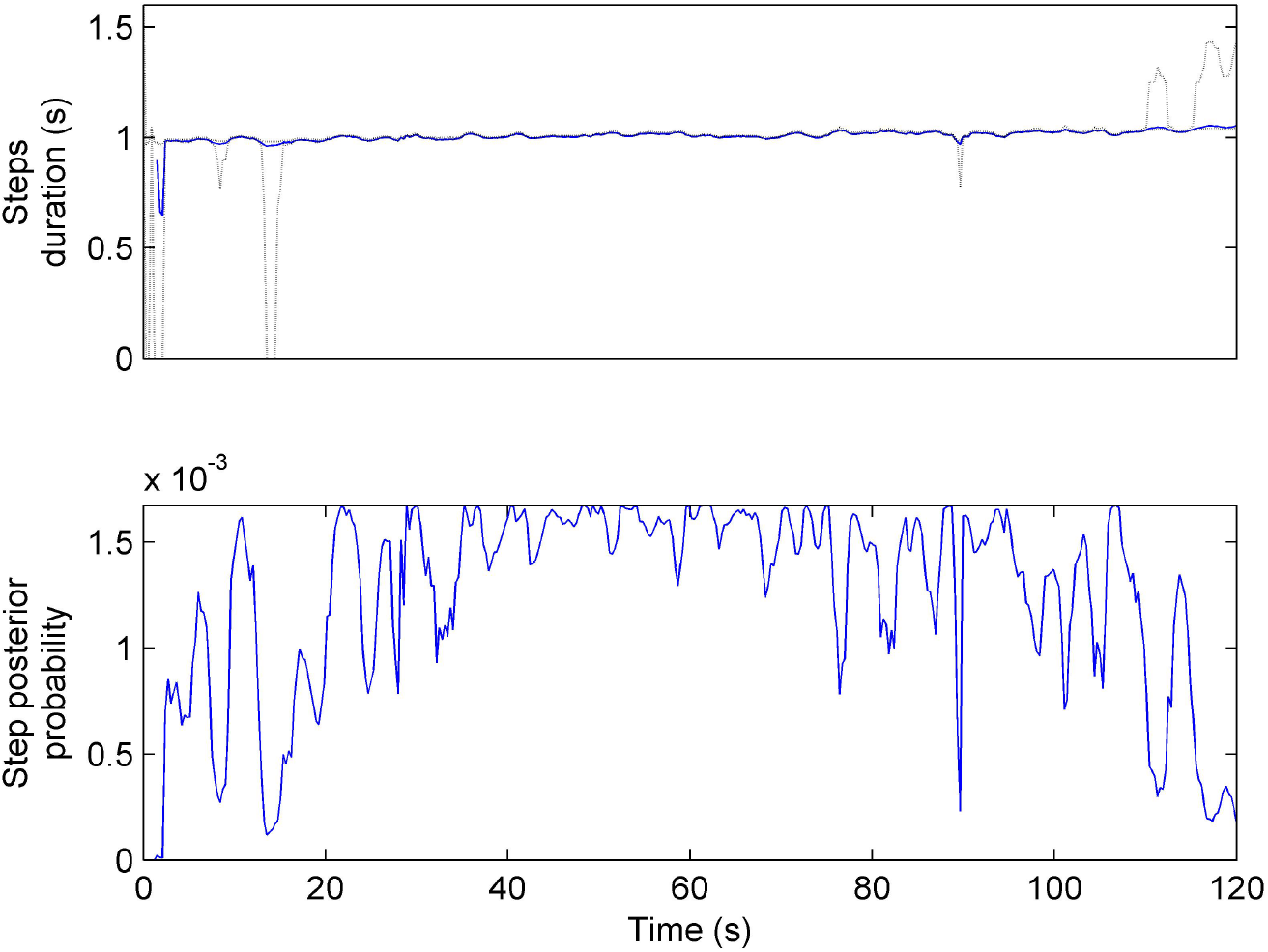
Step durations (top, blue line) and associated posterior probabilities (bottom, blue line) estimated for the same example walk test from experiment #4 as shown in Figure 3. Dotted lines indicate the 95% confidence interval for the estimated step durations.

**Figure 6:**
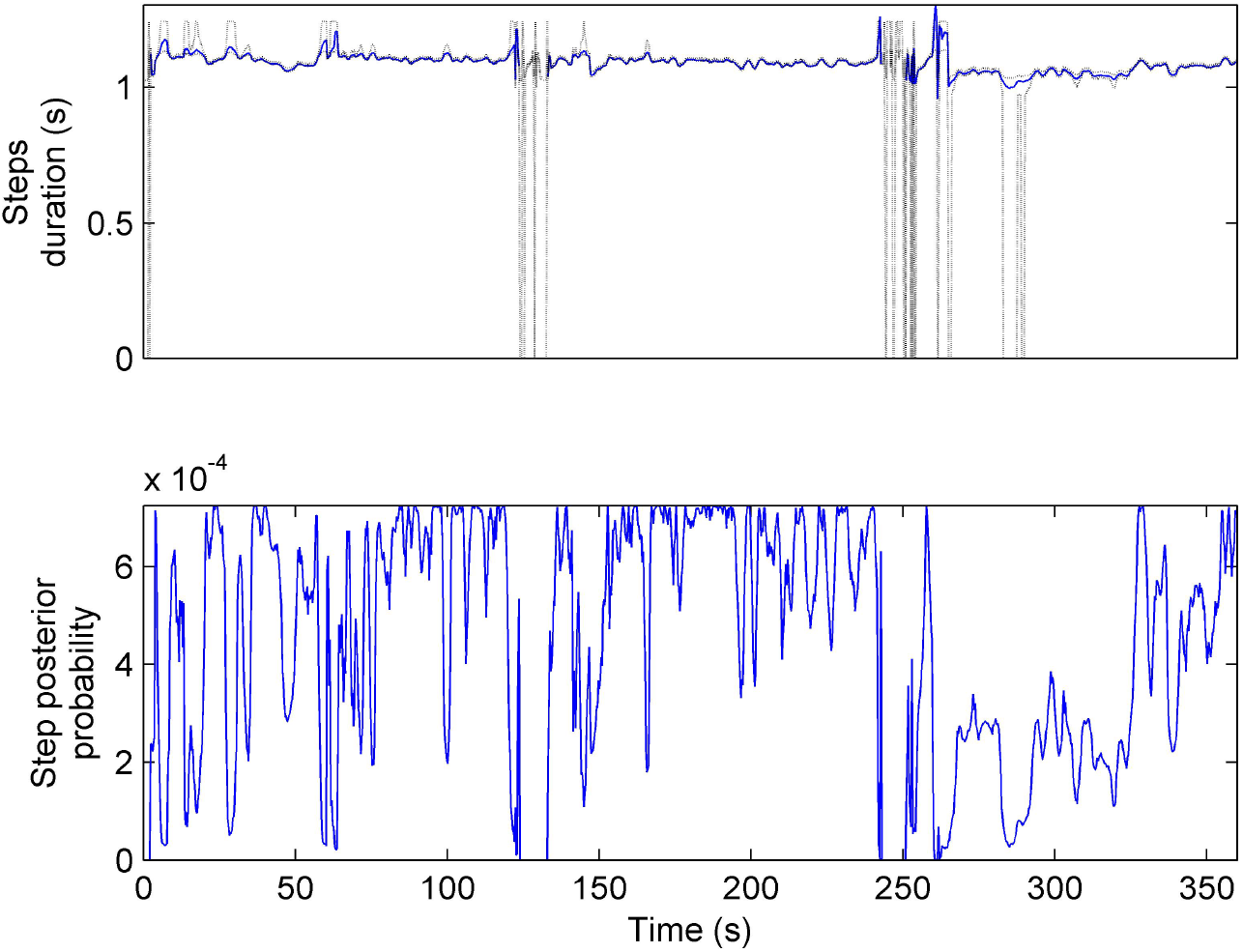
Step durations (top, blue line) and associated posterior probabilities (bottom, blue line) estimated for the same example walk test from experiment #3 as shown in Figure 4. Dotted lines indicate the 95% confidence interval for the estimated step durations.

**Figure 7:**
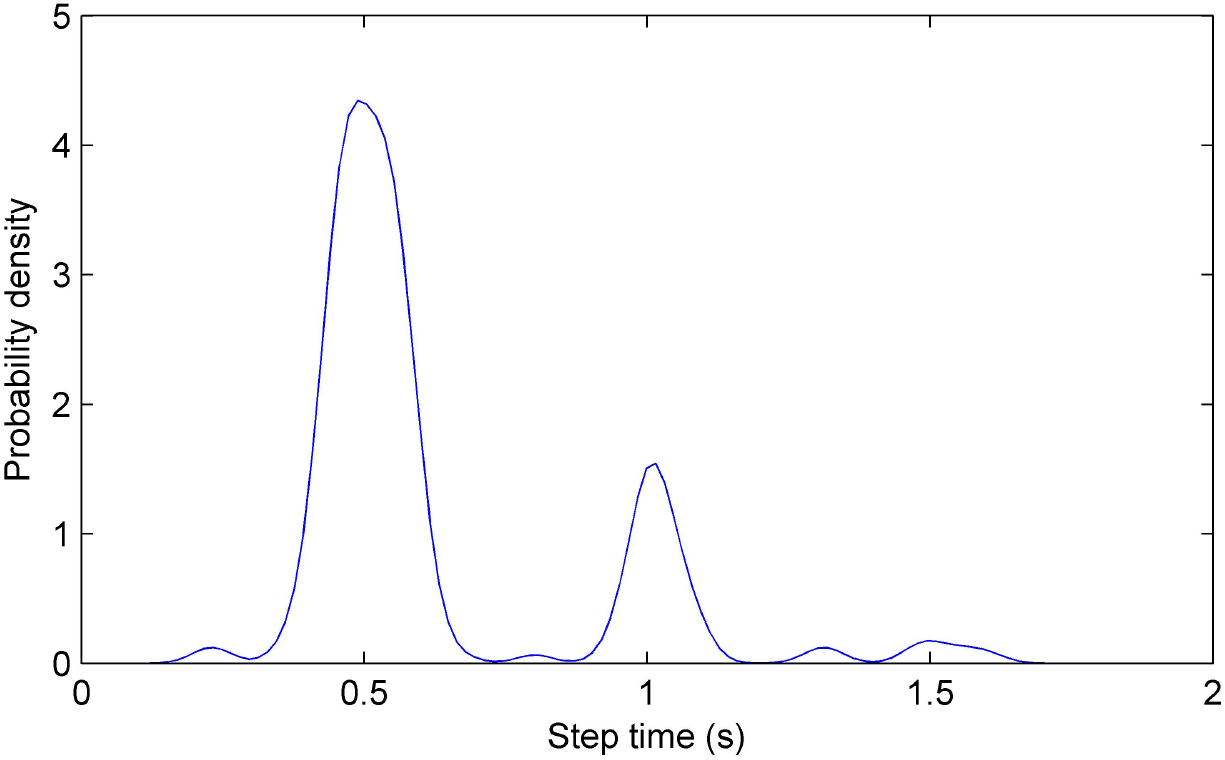
Distribution of step event times τ_step_, Gaussian kernel density estimate for the same example walk test from experiment #4 as shown in Figure 3.

**Figure 8:**
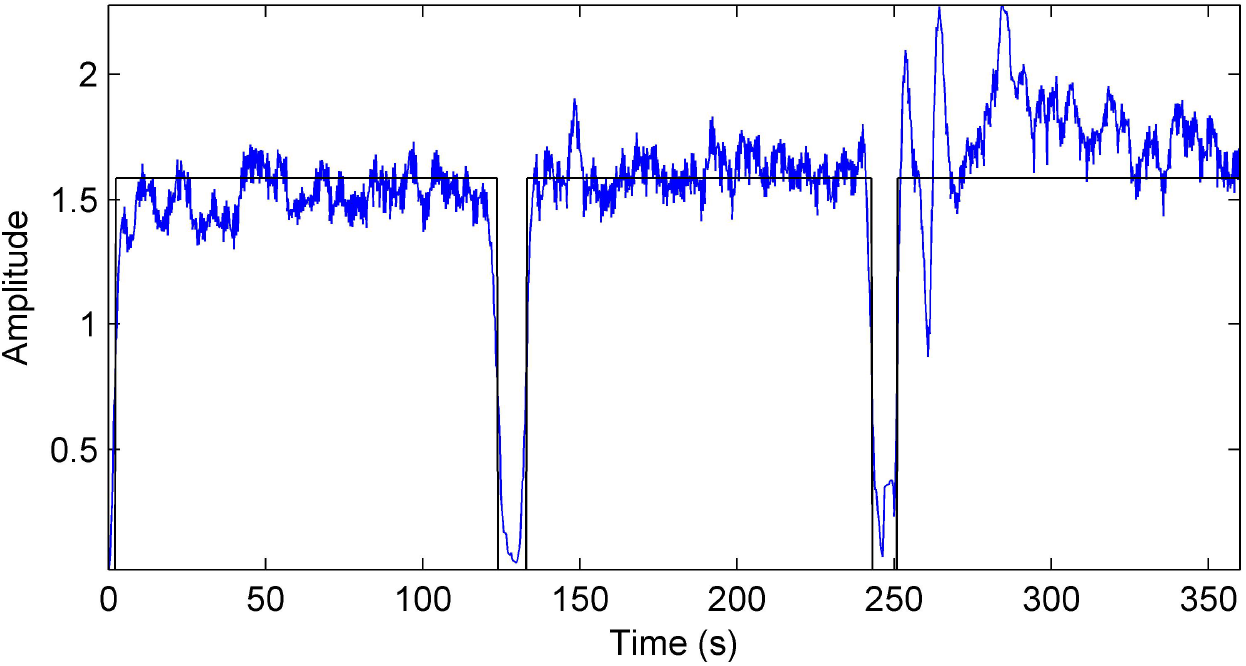
Dynamic acceleration magnitude (blue line) and estimated walk/rest state (black line) for the same example walk test from experiment #4 as shown in Figure 3. Low amplitude indicates rest, high amplitude walk.

We also illustrate the unavoidable ambiguity in estimating *p*_steplike_ (*τ*) induced by high levels of left/right symmetry (Figure 9).

**Figure 9:**
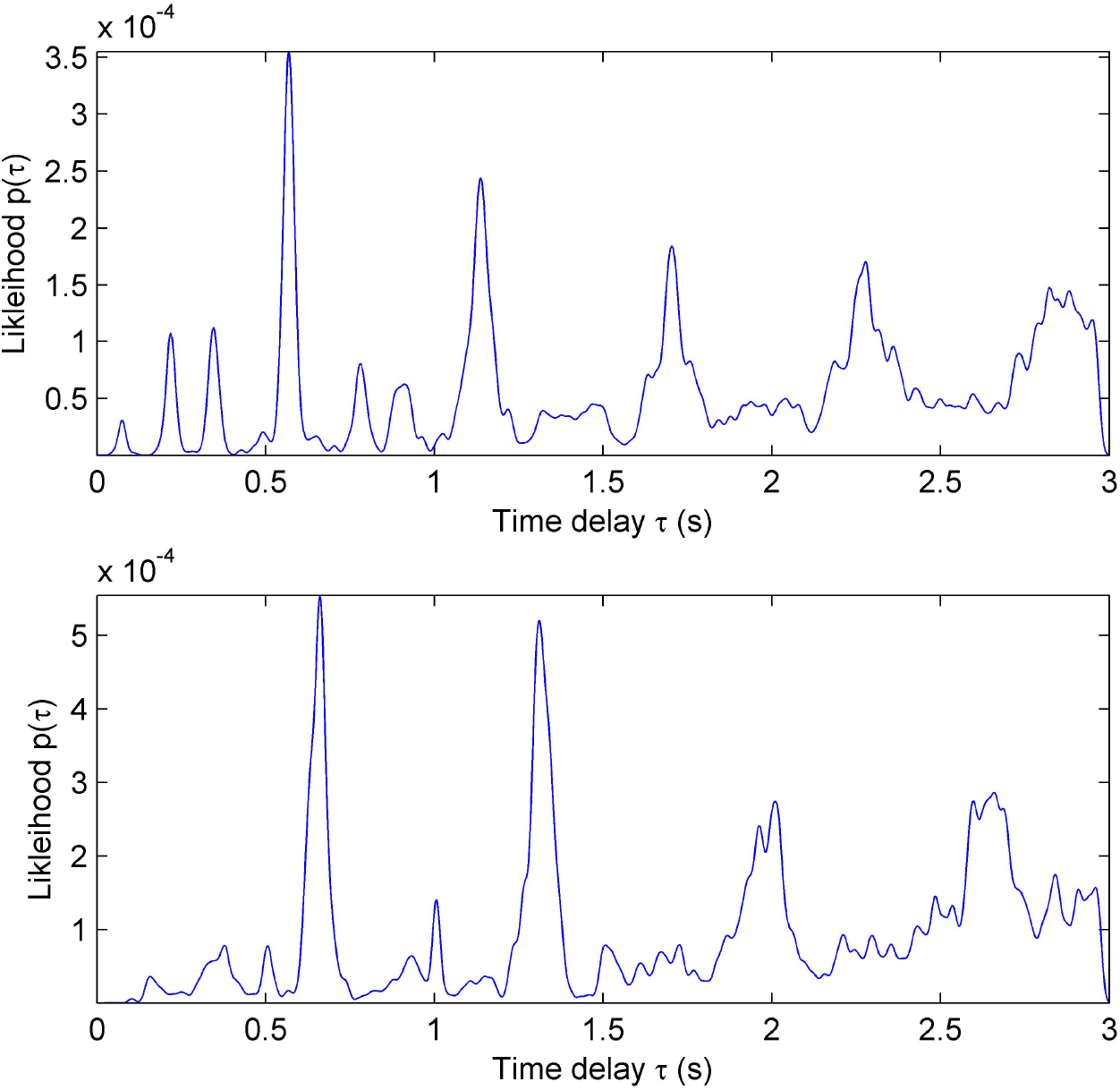
Distribution of step rate likelihoods p_steplike_ (τ), illustrating both asymmetric (top panel) and symmetric (bottom panel) cases. In the asymmetric case, there is a clear distinction between the probability at the fundamental period τ_0_ (largest peak) and the first harmonic 2τ_0_ (second largest peak). Whereas, in the symmetric case, there is little distinction between the two peaks, making it difficult to select the fundamental period without making additional assumptions.

For GPS data from experiment #7, we illustrate an example of estimated walking speed (Figure 10).

**Figure 10:**
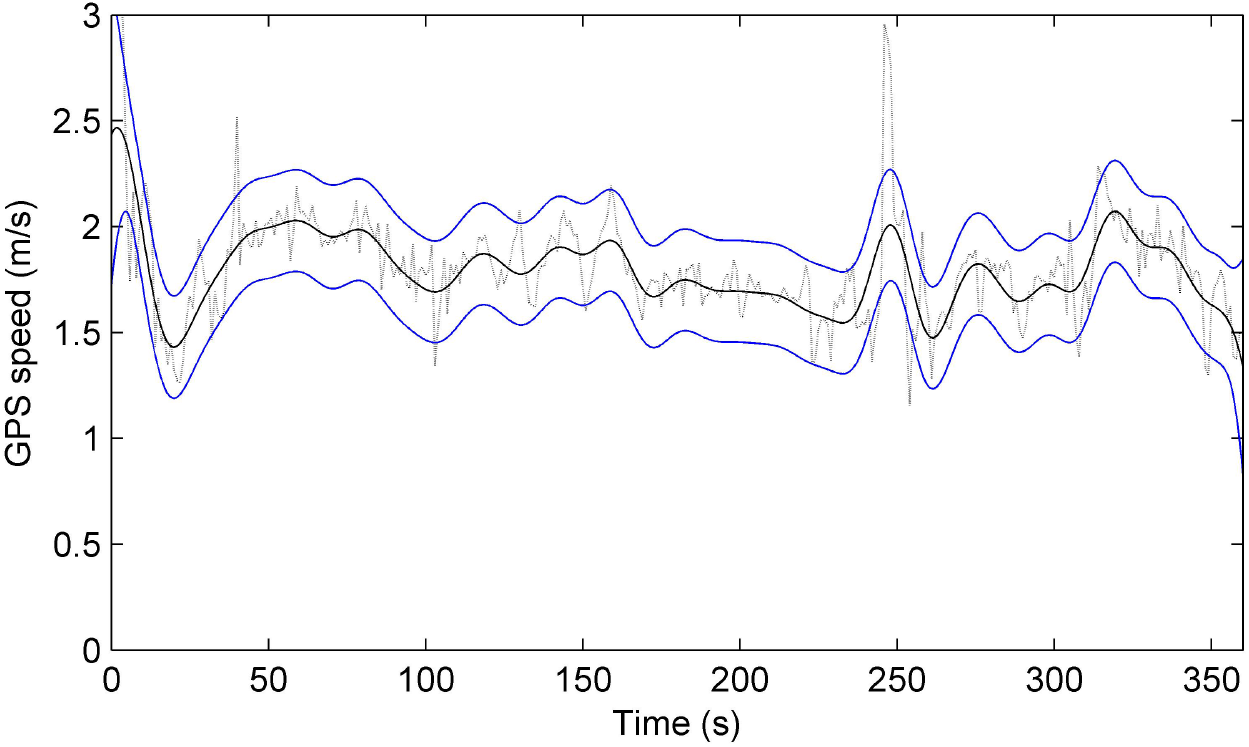
Walking speed along the walking path of a walk test, estimated from GPS position differences and smoothed using Gaussian process estimation, see 30. Most probable (MAP) speed μ_speed_ (t) (black line) is enclosed by ±2 standard deviations (blue lines) computed from the underlying GPS position differences (black dotted line). Example signal from walking experiment #7 conducted on a flat, hard surface at average pace.

### 5.2 Step rate estimation error

For the 59 accelerometer time series in experiments #1-6, the step rate, computed as the inverse step duration 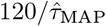, had an estimation error of 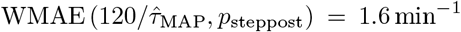 and 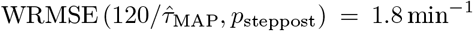, percentage errors 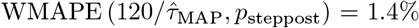 and 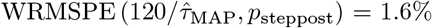. It was found that for experiment #4 the shoulder phone location lead to one very poor outlier estimate with non-negligible posterior probability; therefore, for all three shoulder-located tests the parameters for the step rate prior *p*_stepprior_ (*τ*) in (21) were fixed at mean *μ* = 1.0, standard deviation *σ* = 0.25 and sharpness *p* = 10. Without the fixed prior, the single outlier caused the total estimation error to increase to 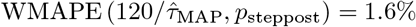. The Bland-Altman agreement plot for these estimates is shown in Figure 11.

**Figure 11:**
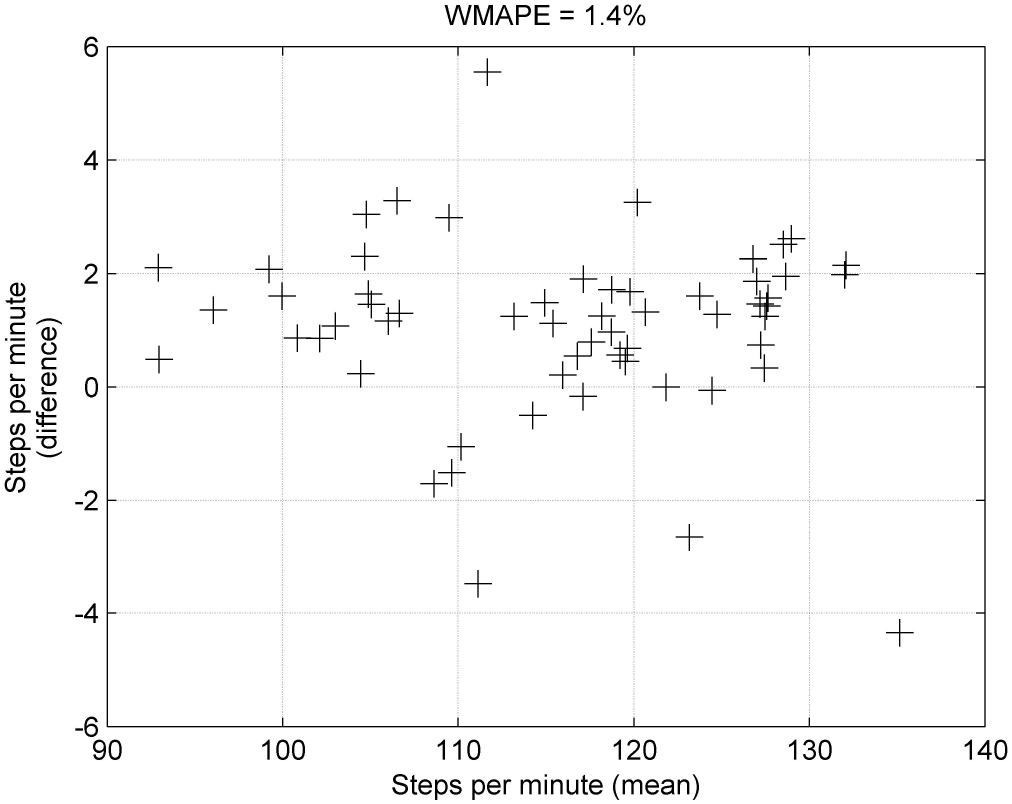
Bland-Altman agreement plot for accelerometer-derived step rate estimation for the data collected in experiments #1-6.

### 5.3 Distance and speed estimation error

Using the 16 GPS traces in experiment #7, the (non-weighted) distance estimation errors were 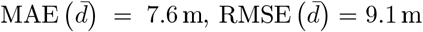 with corresponding percentages 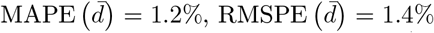. Similarly, for mean walking speed estimates over the whole trace, the estimation errors were 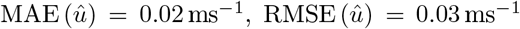 with percentages 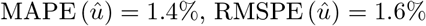. Distance and speed Bland-Altman plots are shown in Figure 12.

**Figure 12:**
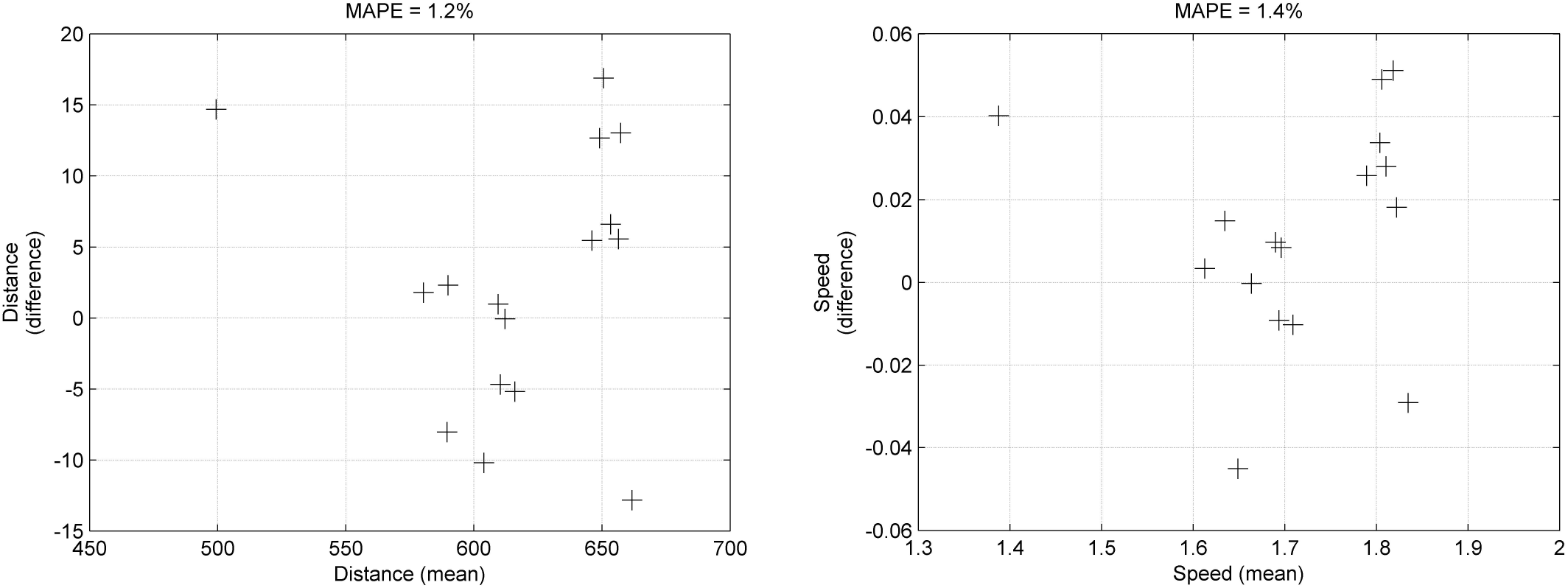
Bland-Altman agreement plots for GPS-derived quantities for the data collected in experiment #7. (Left) total distance walked over each test, and (right) mean walking speed over each test.

## 6 Discussion

Regarding errors, step rates are estimated to within 2% of the reference measurements. While clearly not as accurate as constrained gait lab measurements, this level of error is well within the useful range for reliable measurement of performance outcomes in many non-specialist clinical applications.

For step rates, this is achieved across various terrain, a range of walking speeds and phone locations, including when the phone is held in the hand rather directly mechanically coupled to somewhere on the trunk. As shown in Figure 7, the step event distribution can have significant uncertainty and still provide useful information to “guide” the step rate estimation towards the correct value.

Nonetheless, significant deviations from the protocol can lead to failure which cannot be detected by examining posterior probabilities for signs of low probability, which might otherwise indicate significant measurement uncertainty. In particular, when the phone is worn on the shoulder, sometimes the step events are incorrectly located in such a way which, unfortunately, coincides with incorrect step rate correlation estimates. As a result, these errors combine to lead to very substantial estimation error. This failure was circumvented by choosing an inflexible prior in place of step events, but this “fix” introduces yet another restrictive assumption which imposes constraints on the range of outputs the algorithm can produce. This illustrates one way in which exceeding the limits of the assumptions underlying the derivation of the algorithms can lead to unanticipated failures. It should also be noted that for serious gait impairments, there is no guarantee that these assumptions will hold and the very limited experiments performed here cannot probe this kind of failure.

For GPS-derived measurements, speeds and distances are also estimated to within 2% of the reference measurements. Again, this is accurate enough for non-specialist applications but would not match measurement with higher precision devices. This level of error is achieved for all outdoor walking conditions but it should be stressed that this is only possible with good satellite lock. That means the GPS device must have line-of-sight access to sufficient number of satellites and without this, estimation error is likely to be much worse. For example, the experiments did not test whether such GPS-derived measurements would work in “urban canyons” such as built-up city areas and this remains to be tested. However, the software can provide a measure of sufficient/insufficient GPS satellite lock which may be useful in mitigating this challenge.

## 7 Summary and conclusions

In this paper, we have described the construction of a smartphone-based system implementing a short walk test and using the sensors in the smartphone, developed signal processing algorithms to measure some key performance parameters of basic gait behaviour during the test. We demonstrated using a small number of participants the reliability of these algorithms to some variations within the short walk test protocol. This system could form the basis of a more rigorously tested approach to the measurement of many clinically-relevant aspects of gait behaviour using consumer smartphones, which is a key ambition in modern digital health.

As discussed in the introduction, we emphasise that this report does not in any way claim to be the *definitive* way in which such a system could be devised and implemented. Indeed, as cited above, there are many existing approaches reported in the literature. Nevertheless, we believe that this report illustrates most of the problems which must be addressed in re-purposing consumer smartphones for such an application. For example, the common problems encounter will involve: programming the app to capture sensor data reliably and consistently, defining the test protocols and pre-processing the sensor data to eliminate unwanted variability accordingly, and finally devising robust algorithms to extract performance measures which are reliable under reasonable (and inevitable) deviations from the protocol.

## Acknowledgements

M.L. partially funded by NIH grant UR-Udall Center, award number P50 NS108676. B.B., S.V., B.S., U.H., F.M. and J.O. funded by Signant Health.

## Notes

### Competing Interest Statement

The authors have declared no competing interest.

## References

R. Badawy, Y.P. Raykov, Luc J.W. Evers, Bastiaan R. Bloem, Marjan J. Faber, Andong Zhan, Kasper Claes, and M.A. Little. Automated quality control for sensor based symptom measurement performed outside the lab. Sensors, 18(4): 1215, 2018.

David R. Bassett, Lindsay P. Toth, Samuel R. LaMunion, and Scott E. Crouter. Step counting: A review of measurement considerations and health-related applications. Sports Medicine, 47(7):1303–1315, 2017. 1179-2035.

Daniel N. Cagney, Joohee Sul, Raymond Y. Huang, Keith L. Ligon, Patrick Y. Wen, and Brian M. Alexander. The FDA NIH Biomarkers, EndpointS, and other Tools (BEST) resource in neuro-oncology. Neuro-Oncology, 20(9):1162–1172, 2017.

Michelle Crouthamel, Emilia Quattrocchi, Sarah Watts, Sherry Wang, Pamela Berry, Luis Garcia-Gancedo, Valentin Hamy, and Rachel E. Williams. Using a ResearchKit smartphone app to collect rheumatoid arthritis symptoms from real-world participants: Feasibility study. JMIR Mhealth Uhealth, 6(9):e177, 2018. 13.9.2018.

J. A. DiMasi, H.G. Grabowski, and R.W. Hansen. Innovation in the pharmaceutical industry: New estimates of R&D costs. Journal of Health Economics, 47:20–33, 2016.

V. Hamy, L. Garcia-Gancedo, A. Pollard, A. Myatt, J. Liu, A. Howland, P. Beineke, E. Quattrocchi, R. Williams, and M. Crouthamel. Developing smartphone-based objective assessments of physical function in rheumatoid arthritis patients: The PARADE study. Digital Biomarkers, 4:26–43, 2020.

K. I. Kaitin and J.A. DiMasi. Pharmaceutical innovation in the 21st century: New drug approvals in the first decade, 2000-2009. Clinical Pharmacology and Therapeutics, 89(2):183–188, 2011.

C.F.F. Karney. Transverse Mercator with an accuracy of a few nanometers. Journal of Geodesy, 85(8):475–485, 2011.

S.-J. Kim, K. Koh, S. Boyd, and D. Gorinevsky. l1 trend filtering. SIAM Review, 51(2):339–360, 2009.

Florian Lipsmeier, Kirsten I. Taylor, Timothy Kilchenmann, Detlef Wolf, Alf Scotland, Jens Schjodt-Eriksen, Wei-Yi Cheng, Ignacio Fernandez-Garcia, Juliane Siebourg-Polster, Liping Jin, Jay Soto, Lynne Verselis, Frank Boess, Martin Koller, Michael Grundman, Andreas U. Monsch, Ronald B. Postuma, Anirvan Ghosh, Thomas Kremer, Christian Czech, Christian Gossens, and Michael Lindemann. Evaluation of smartphone-based testing to generate exploratory outcome measures in a phase 1 parkinson’s disease clinical trial. Movement Disorders, 33(8):1287–1297, 2018.

M.A. Little. Machine Learning for Signal Processing. Oxford University Press, 2019.

M.A. Little and N.S. Jones. Generalized methods and solvers for noise removal from piecewise constant signals. I. background theory. Proceedings of the Royal Society A: Mathematical, Physical and Engineering Science, 467(2135), 2011.

J.N.S. Matthews. Introduction to Randomized Controlled Clinical Trials. CRC Press, 2nd edition, 2006.

Michael V. McConnell, Anna Shcherbina, Aleksandra Pavlovic, Julian R. Homburger, Rachel L. Goldfeder, Daryl Waggot, Mildred K. Cho, Mary E. Rosenberger, William L. Haskell, Jonathan Myers, Mary Ann Champagne, Emmanuel Mignot, Martin Landray, Lionel Tarassenko, Robert A. Harrington, Alan C. Yeung, and Euan A. Ashley. Feasibility of obtaining measures of lifestyle from a smartphone app: The myheart counts cardiovascular health study. JAMA Cardiology, 2(1): 67–76, 2017.

A. Middelweerd, J.S. Mollee, C.N. van der Wal, J. Brug, and S.J. te Velde. Apps to promote physical activity among adults: A review and content analysis. International Journal of Behavioral Nutrition and Physical Activity, pages 1–9, 2014.

D. Mohammadi. ResearchKit: a clever tool to gather clinical data. The Pharmaceutical Journal, 2015.

J. Pearl. Causality: Models, Reasoning and Interference. Cambridge University Press, Cambridge, UK, 2009.

W. H. Press, S.A. Teukolsky, W.T. Vetterling, and B.P. Flannery. Numerical Recipes in C: The Art of Scientific Computing. Cambridge University Press, Cambridge, UK, 2002.

L. Shi, J.K. Nielsen, J.R. Jensen, M.A. Little, and M.G. Christensen. Robust bayesian pitch tracking based on the harmonic model. IEEE/ACM Transactions on Audio, Speech, and Language Processing, 27(11):1737–1751, 2019.

X. Sun. A pitch determination algorithm based on subharmonic-to-harmonic ratio. In Sixth International Conference on Spoken Language Processing (ICSLP 2000), volume 4, pages 676–679, Beijing, China, 2000.

F. Vaverka, M. Elfmark, Z. Svoboda, and M. Janura. System of gait analysis based on ground reaction force assessment. Acta Gymnica, 45(4):187–193, 2015.

